# An efficient genome editing strategy to generate putative null mutants in *Caenorhabditis elegans* using CRISPR/Cas9

**DOI:** 10.1101/391243

**Authors:** Han Wang, Heenam Park, Jonathan Liu, Paul W. Sternberg

## Abstract

Null mutants are essential for analyzing gene function. Here, we describe a simple and efficient method to generate *Caenorhabditis elegans* null mutants using CRISPR/Cas9 and short single stranded DNA oligo repair templates to insert a universal 43-nucleotide-long stop knock-in (STOP-IN) cassette into the early exons of target genes. This cassette has stop codons in all three reading frames and leads to frameshifts, which will generate putative null mutations regardless of the reading frame of the insertion position in exons. The STOP-IN cassette also contains an exogenous Cas9 target site that allows further genome editing and provides a unique sequence that simplifies the identification of successful insertion events via PCR. As a proof of concept, we inserted the STOP-IN cassette right at a Cas9 target site in *aex-2* to generate new putative null alleles by injecting preassembled Cas9 ribonucleoprotein and a short synthetic single stranded DNA repair template containing the STOP-IN cassette and two 35-nucleotide-long homology arms identical to the sequences flanking the Cas9 cut site. We showed that these new *aex-2* alleles phenocopied an existing loss-of-function allele of *aex-2*. We further showed that the new *aex-2* null alleles could be reverted back to the wild-type sequence by targeting exogenous Cas9 cut site included in the STOP-IN cassette and providing a single stranded wild-type DNA repair oligo. We applied our STOP-IN method to generate new putative null mutants for additional 20 genes, including three pharyngeal muscle-specific genes (*clik-1, clik-2*, and *clik-3*), and reported a high insertion rate (46%) based on the animals we screened. We showed that null mutations of *clik-2* cause recessive lethality with a severe pumping defect and *clik-3* null mutants have a mild pumping defect, while *clik-1* is dispensable for pumping. We expect that the knock-in method using the STOP-IN cassette will facilitate the generation of new null mutants to understand gene function in *C. elegans* and other genetic model organisms.

**Summary:** We report a simple and efficient CRISPR/Cas9 genome editing strategy to generate putative null *C. elegans* mutants by inserting a small universal stop knock-in (STOP-IN) cassette with stop codons in three frames and frameshifts. The strategy is cloning-free, with the mixture consisting of preassembled Cas9 ribonucleoprotein and single stranded repair DNA oligos directly injected into gonads of adult *C. elegans*. The universal STOP-IN cassette also contains a unique sequence that simplifies detection of successful knock-in events via PCR and an exogenous Cas9 target sequence that allows further genome editing.

## Materials and Methods

### Maintenance of Strains

All *C. elegans* strains were maintained on NGM plates seeded with the *E. coli* strain OP50 as food, at room temperature, as described (Brenner 1974). The following strains were used: wild type (the N2 Bristol strain), JT3 *aex-2(sa3) X*, and a balancer strain FX30236 *tmC30[ubc-17(tmIs1243)]X* (Dejima *et al*. 2018). Strains generated in this study are listed in Supplemental Table S1.

### Design of guide RNA sequences

All crRNAs and the universal tracrRNA were chemically synthesized (the Alt-R system from IDT, Coralville, IA). Synthetic crRNAs and tracrRNA were dissolved in nuclease-free water (IDT) to make 100 μM working stocks. The guide sequences of all crRNAs used in this study are provided in Supplemental Table S2. We used a CRISPR guide sequence selection tool (http://genome.sfu.ca/crispr/search.html) designed for the *C. elegans* genome (Au *et al*. 2018) to identify guide sequences within the coding sequences of target genes; one guide sequence was selected for each gene of interest. We only selected guide sequences followed by the PAM site (5’-NGG-3’) for wild-type *Streptococcus pyogenes* Cas9. The guide sequences in early exons shared by all known isoforms of the gene of interest were preferred, as the early stop codons and frameshifts introduced by our STOP-IN method in these regions were more likely to produce putative null mutants.

### Design of repair DNA oligos

Single stranded DNA oligos with short homology arms (~ 30-60 nucleotides for each arm) efficiently serve as repair templates to insert up to 130 nucleotides into the *C. elegans* genome with CRISPR (Paix *et al*. 2014, 2017). All single stranded DNA repair oligos were synthesized (4nM, Ultramer DNA oligos from IDT, Coralville, IA) and were diluted in nuclease-free water (IDT). The sequences of DNA repair oligos used in this study are described in Supplementary Table S2. Our universal stop knock-in (STOP-IN) cassette (43 nucleotides, 5’-GGGAAGTTTGTCCAGAGCAG<u>AGG</u>TGACTAAGTGATAAGCTAGC-3’) is composed of an exogenous Cas9 target site with a PAM site (underlined) that has been shown to work in *C. elegans* (Paix *et al*. 2017), multiple stop codons in all three reading frames, and the recognition site of the restriction enzyme *NheI* (Figure 1B).

**Figure 1.**
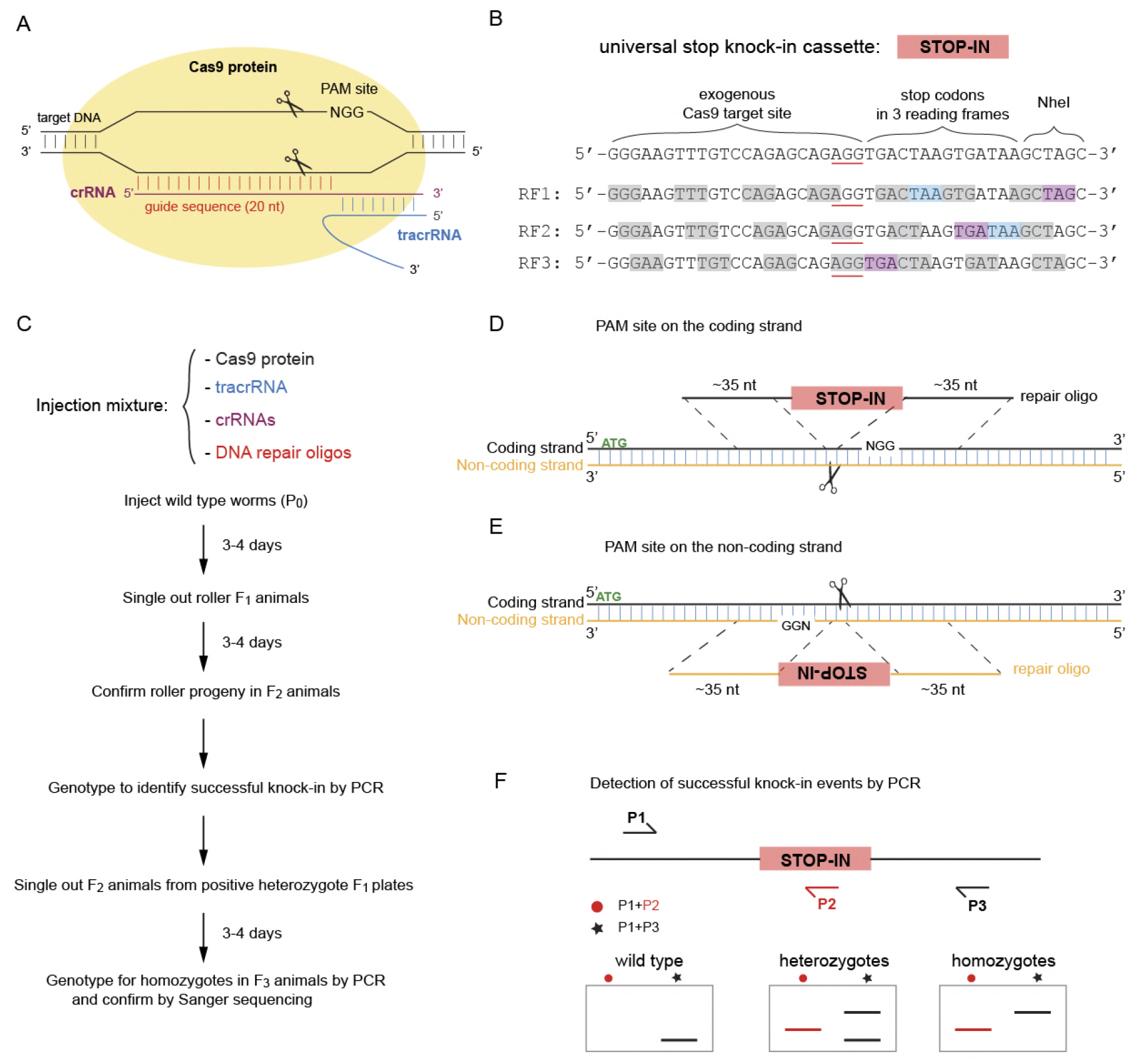
The CRISPR pipeline to generate null mutants by inserting a universal stop knock-in (STOP-IN) cassette. (**A**) Diagram showing the recognition of the target DNA by Cas9 ribonucleoprotein from *Streptococcus pyogenes*. The ribonucleoprotein complex of Cas9 with crRNA and tracrRNA recognizes the target DNA with the first 20 nucleotides of crRNA homology (the guide sequence) preceding a PAM site (‘5-NGG-3’) and generates a double-strand break at 3 nucleotides 5’ of the PAM site. (**B**) Design of the 43-nucleotide-long universal stop knock-in (STOP-IN) cassette (represented by the red rectangle) used in this study, which includes an exogenous Cas9 target site (the PAM site is underlined), stop codons and frameshifts in three frames, and the recognition site for the restriction enzyme NheI. RF, reading frame, as indicated by grey boxes. Cyan and purple boxes represent the stop codon TAA and TGA, respectively. Reading frames 1 and 2 have two stop codons; reading frame 3 has one stop codon. (**C**) The recipe of the ribonucleoprotein injection mixture and the workflow of the CRISPR protocol to generate null mutants. With the co-conversion method, two crRNAs and two DNA repair oligos (one pair for the co-conversion marker, *dpy-10 or unc-58*, and the other pair for the target gene) are included in the same injection mixture; the mixture is directly injected into the gonads of adult hermaphrodites. Animals with genomes affected by CRISPR/Cas9 and template-directed repair are identified by the co-conversion marker phenotype; in the figure, these are roller animals with *dpy-10* conversion. (**D and E**) Design of the DNA repair oligo for the target gene. The oligo contains two ~ 35-nucleotide-long homology arms identical to the sequences flanking the Cas9 cut site and in the same strand as the PAM site (5’-NGG-3’). If the PAM site is on the coding strand (1D), the STOP-IN cassette sequence as shown in (1B) is included in between the two homology arms to comprise the DNA repair oligo. If the PAM site is on the non-coding strand (1E), the reverse complement of the STOP-IN cassette sequence as shown in (1B) is added between the two homology arms to make the DNA repair oligo. (**F**) Diagram showing the placement of primers for PCR reactions to detect successful knock-in events. P1 and P3 are outer primers that are specific to the target gene, while P2 is a common internal primer within the universal STOP-IN cassette. These primers can be used to detect the successful insertion of the STOP-IN cassette via PCR, as shown in the simulated agarose gels below.

To achieve high knock-in efficiency, we implemented two rules for the design the single-strand DNA repair oligos. First, the efficiency of HDR correlates inversely with the distance between homology arms and the Cas9 cut site (Paquet *et al*. 2016). Thus, each single stranded DNA repair oligo contains two unique homology arms (~35 nucleotides for each) that are identical to sequences immediately flanking the Cas9 cut site, such that the universal STOP-IN cassette will be directly inserted to the cut site. In addition, successful insertion will disrupt the original Cas9 target site preventing the knock-in locus from being re-cut by Cas9, and removing the need to introduce additional mutations in the repair template. Second, it has also been demonstrated that the sense of the single stranded DNA repair oligo relative to the PAM site affects the genome editing efficiency: if the modification occurs at 5’ to the PAM site (5’-NGG-3’), the DNA repair oligo containing sequence from the same strand as the PAM site has higher knock-in efficiency; if the modification occurs at 3’ to the PAM site, the DNA repair oligo containing the reverse complement sequence of the strand with the PAM site has higher knock-in efficiency (Barbara J. Meyer, personal communication). As the universal STOP-IN cassette needs to be inserted at the Cas9 cut site in the coding strand (3 nt at the 5’ end of the PAM site) to generate putative null mutants, the single stranded DNA oligo should be in the same strand that carries the PAM (5’-NGG-3’). Thus, each single stranded DNA repair oligo (~113 nucleotides) contains the STOP-IN cassette and two unique homology arms (~35 nucleotides for each) identical to the flanking sequences of the corresponding Cas9 cut site. The direction of the universal STOP-IN cassette (sense or anti-sense) in the DNA repair oligo, is dependent on whether the PAM site is in on the coding strand or the non-coding strand (Figure 1E), such that the stop codons will be inserted into the coding sequence of the target gene at the Cas9 cut site via HDR. A detailed step-by-step protocol to design DNA repair oligos is included in Supplemental File S1.

### Microinjection of Cas9 ribonucleoprotein

We used the co-conversion method (Arribere *et al*. 2014) for all the CRISPR/Cas9 genome editing experiments in this study. We injected preassembled Cas9 ribonucleoprotein and synthetic single stranded DNA oligos as repair template (Paix *et al*. 2015; Figure 1C). Using the plasmid pHO4d-Cas9 (Addgene plasmid # 67881), which was a gift from Michael Nonet (Hua Fu *et al*. 2014), the Cas9 protein (eluted in the buffer with 20 mM Hepes pH7.5, 150 mM KCl, 10% Glycerol, 1 mM DTT) was made as previously described (Paix *et al*. 2015). To anneal tracrRNA and crRNAs at 1:1 ratio, 2.5 μl of 100 μM tracrRNA, 0.5 μl of 100 μM *dpy-10* crRNA (for co-conversion), and 2.0 μl of 100 μM crRNA targeting the gene of interest were mixed in a PCR tube, incubated at 94 °C for 2 min in a PCR machine, and cooled down to room temperature. The injection solution was prepared as follows: 3.4 μg/μL Cas9 protein, 45.9 μM the annealed guide RNA mixture, 0.51 μM DNA repair oligo for the gene of interest, and 0.25 μM DNA repair oligo for the co-conversion marker *dpy-10*. The solution was spun at 14549 rcf for 1 min, put on ice and can be used right away. The mixture was injected into the germline syncytium of wild type *C. elegans* according to a standard microinjection procedure (Mello and Fire 1995). We injected 15-20 young adults (P0), cultured each injected P0 onto its own NGM Petri plate at room temperature, and picked F_1_ progeny with the co-conversion phenotype after 3-4 days. For the experiment to obtain *aex-2* revertants, the *aex-2(sy1078)* animals were used for injection. When using *unc-58* as the co-conversion marker, we slightly modified the protocol by changing the ratio of crRNAs in the annealed guide RNA mixture (1.0 μl of 100 μM *unc-58* crRNA, and 1.5 μl of 100 μM crRNA for the target gene) and the final concentration of the repair template for *unc-58* (0.51 μM) in the injection solution. A step-by-step protocol is provided in Supplemental File S1.

### Screen for successful knock-in events by PCR

3-4 days after microinjection, we selected the F_1_ animals with the dominant co-conversion marker phenotype (roller phenotype for *dpy-10*), singled them onto individual NGM Petri plates, and confirmed the presence of the co-conversion marker phenotype in F_2_ progeny (Figure 1C). As CRISPR/Cas9 via injecting preassembled ribonucleoprotein may induce targeted somatic mutations in the F_1_ progeny (Cho *et al*. 2013), we only genotyped F_2_ progeny to identify F_1_ animals with heritable genome editing. For each of those F_1_ animals that produced F_2_ progeny with the co-conversion marker, indicating heritable genome editing occurred, about 10 young F_2_ animals were lysed and genotyped (See Supplemental File S1 for a detailed protocol). Two PCR reactions were used: one using the common reverse inner primer (oHP013r: 5’-GCTTATCACTTAGTCACCTCTGCTC-3’) within the repair template and a forward primer specific to the target gene to identify F_1_ animals with successful insertion of the STOP-IN cassette; the other one using two outer primers specific to the target gene to determine if the F_1_ animals were heterozygotes for the knock-in allele (Figure 1F). The two outer primers specific to the target gene were chosen to get a relative small PCR amplicon (< 500 bp) such that a small size difference (43 bp) can easily be detected on a 2.5% agarose gel. For positive F_1_ animals that were heterozygotes, several F_2_ progeny from each plate were singled out to identify homozygote F_2_ animals by genotyping F_3_ progeny using the same outer primer set. The successful insertion of the STOP-IN cassette in the homozygous animals was confirmed by Sanger sequencing (Laragen, Culver City, CA) of PCR products with the two outer primers. Primer sequences used for genotyping is provided in Supplemental Table S2. The original F_1_ animals in the F_1_ plates that had predicted knock-in bands from both PCR reactions were considered to be positive F_1_ clones with the desired knock-in alleles. The knock-in efficiency (“% of KI” in Table 1) for each genome editing experiment was calculated as the number of positive F_1_ clones divided by the total number of F_1_ plates genotyped.

**Table 1.**
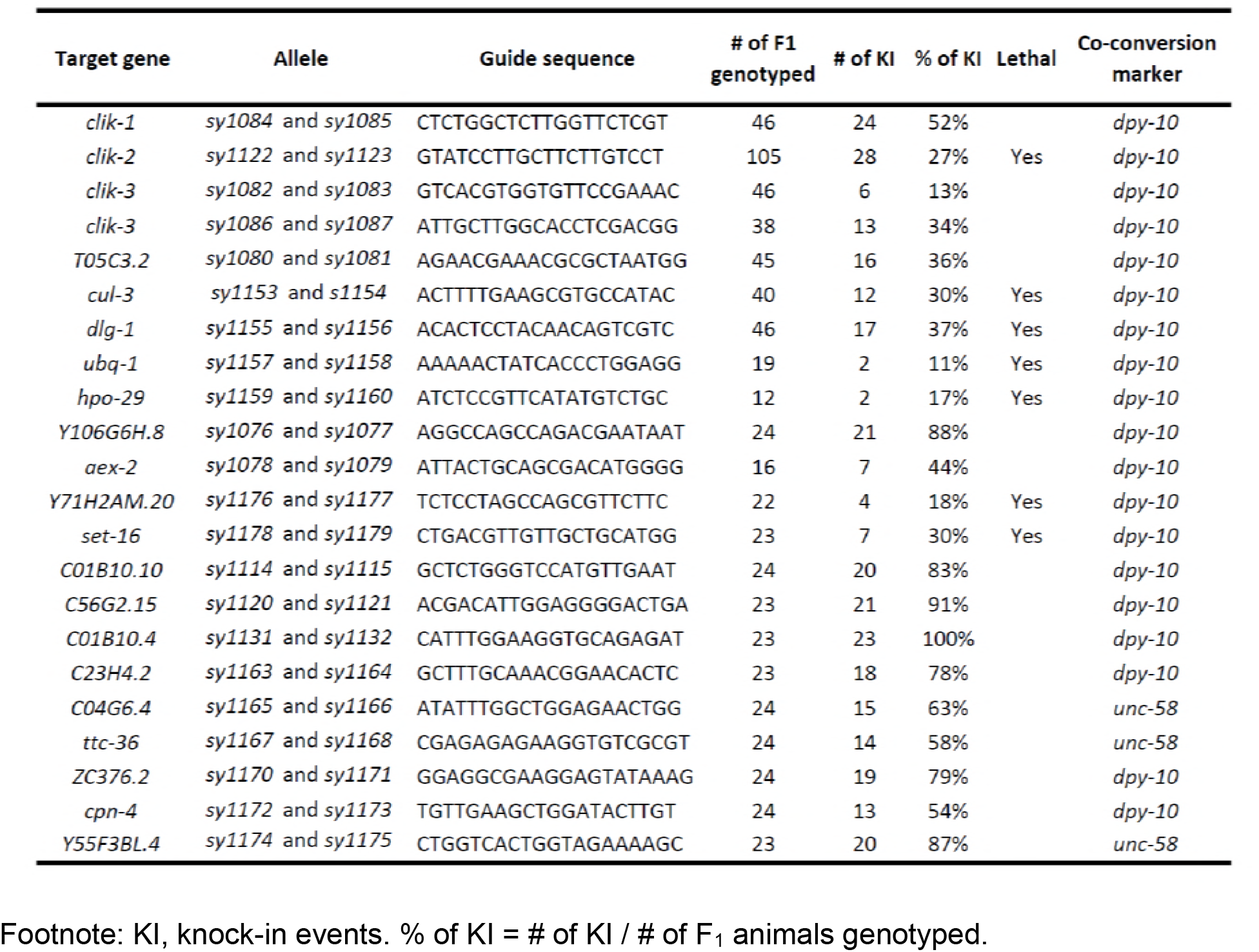
Summary of our CRISPR/Cas9 knock-in results.

For reversion of the *aex-2(sy1078)* null mutants, we screened the F_1_ generation for non-constipated in F_1_ animals using a dissecting microscope and transferred each of candidates onto individual plates. Several wild-type F_2_ progeny from each plate were singled out to identify homozygotes for revertant *aex-2* alleles by the absence of constipated worms in the next generation. Likely homozygous *aex-2* revertants were genotyped with the same set of outer primers used to detect *aex-2(sy1078)* and the revertant wild-type alleles were confirmed by Sanger sequencing.

### Behavioral assays

The defecation motor program was assayed as previously described (Wang *et al*. 2013) with modifications. Briefly, L4 animals were collected, and then 24 hours later transferred to new NGM Petri plates seeded with *E. coli* OP50, and observed under a dissecting stereomicroscope. Each worm was scored for six consecutive defecation cycles. The posterior body wall muscle contraction (pBoc) and enteric muscle contraction (EMC) during each defecation cycle were recorded using Etho, an event logging Java program (http://depts.washington.edu/jtlab/software/otherSoftware.html) (Thomas 1990). “EMC per cycle” for each animal was calculated as the ratio of EMC to pBoc. Ten individual animals were assayed for each genotype.

Pumping assays were performed as follows: 24 hours prior to the assay, L4 animals of each genotype were moved onto fresh NGM Petri plates with *E. coli* OP50. The next day, adult animals were picked in pairs onto new NGM plates seeded with OP50 and allowed to acclimate for 10 minutes. Animals were then recorded on a Wild Makroskop M420 dissecting microscope at 80x magnification for 30 seconds. The number of pumping events observed in 30 seconds was doubled to obtain the pumping rate (pumps/min).

### Microscopy

DIC images of the constipated worms in Figure 2D were taken using a Plan Apochromat 40x/1.4 Oil DIC objective in a Zeiss Imager Z2 equipped with an Axiocam 506 Mono camera using ZEN Blue 2.3 software.

**Figure 2.**
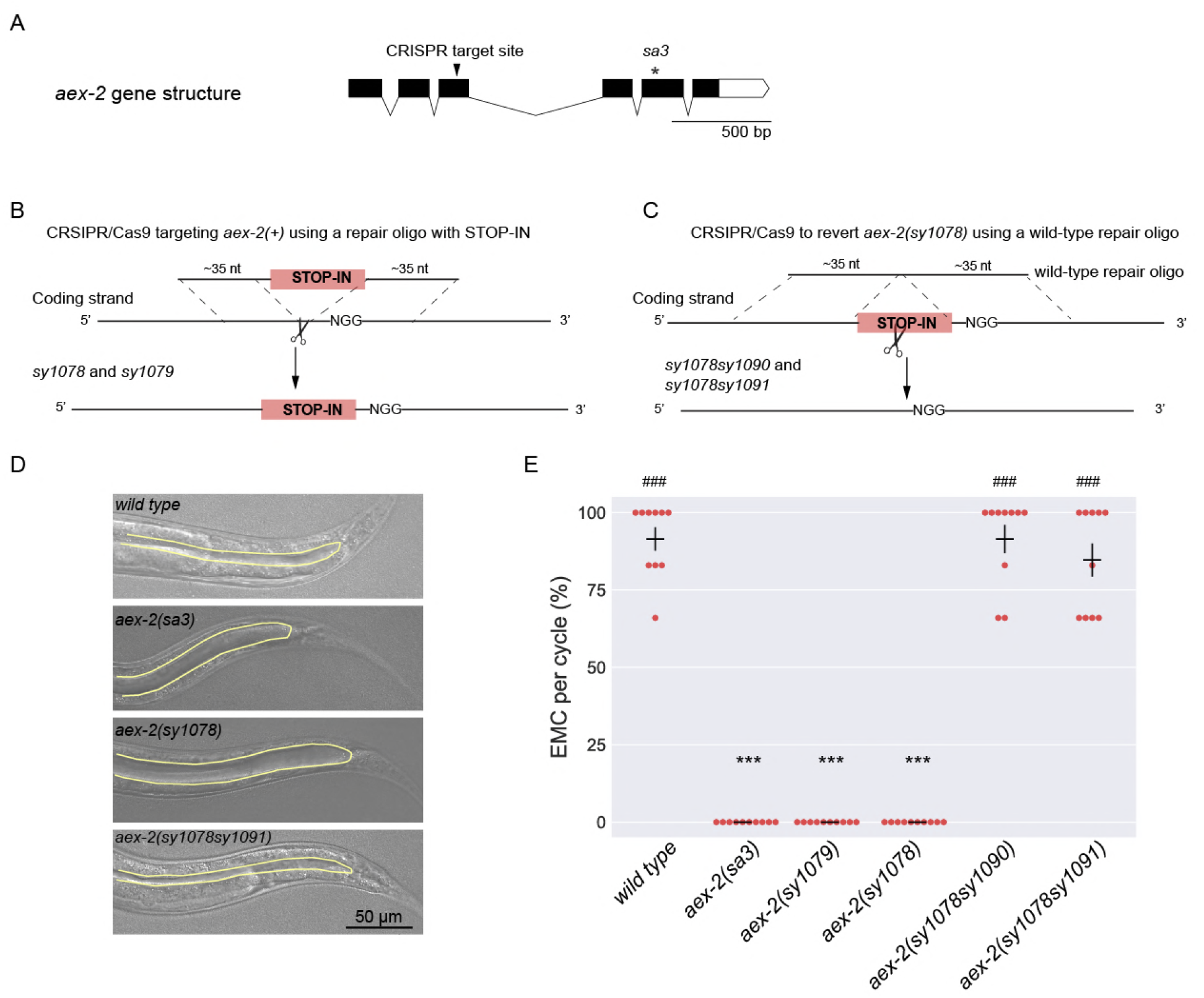
Genome engineering of the *aex-2* gene. (**A**) The gene structure of *aex-2*. The black arrowhead indicates the position of the Cas9 target site in the third exon of *aex-2*, which we used to insert the universal STOP-IN cassette to generate the two new *aex-2* null alleles, *sy1078* and *sy1079*. The asterisk denotes the mutation sa3, a strong loss-of-function allele of *aex-2*. Black boxes are exons and white box is the 3’ UTR. The lines connecting the black boxes are introns. Scale bar, 500 bp. (**B**) Scheme showing the generation of putative *aex-2* null alleles, *sy1078* and *sy1079*, using CRISPR/Cas9 and a DNA repair oligo with the universal STOP-IN cassette. (**C**) Diagram showing the reversion of *aex-2(sy1078)* to wild-type sequence, using the exogenous Cas9 target site included in the STOP-IN cassette and a wild-type DNA oligo with two ~ 35-nucleotide-long homology arms. (**D**) Differential Interference Contrast (DIC) images showing the constipation phenotypes in young adults with indicated genotypes. Yellow lines outline the posterior intestinal lumens. Compared to wild-type and *aex-2(sy1078sy1091)* animals, *aex-2(sa3)* and *aex-2(sy1078)* mutants both have distended intestinal lumens. Scale bar, 50 μm. (**E**) Quantification of enteric muscle contractions (EMC) during the defecation motor program in adult animals with indicated genotypes. Each dot represents the percentage of EMC observed per defecation cycle from an individual animal. n= 10 animals for all genotypes. Horizontal bars indicate the mean; vertical bars are SEM. ***, P < 0.001 and **, P < 0.01, compared to wild type; ###, P < 0.001 and ##, P < 0.01, compared to *aex-2(sy1078)*; One-way ANOVA with Dunn’s multiple comparisons test.

### Statistics

Where appropriate, Chi-square test or One-way ANOVA with Tukey’s post-test or with Dunn’s multiple comparisons test were applied (GraphPad Prism) to determine if there were significant differences between genotypes, as indicated in results or figure legend.

### Statistics

Where appropriate, Chi-square test or One-way ANOVA with Tukey’s post-test or with Dunn’s multiple comparisons test were applied (GraphPad Prism) to determine if there were significant differences between genotypes, as indicated in results or figure legend.

### Data Availability Statement

Strains and plasmids are available upon request. The authors affirm that all data necessary for confirming the conclusions of the article are present within the article, figures, tables, and supplemental information.

## Introduction

An essential goal of genetics is to understand gene function. Loss-of-function or null mutants, in which gene activity is reduced or completely lost, are valuable reagents that enable this goal. *Caenorhabditis elegans* has become an important model organism since the first description of its genetics by Brenner (Brenner 1974), allowing study of conserved mechanisms underlying many important biological processes, such as programed cell death, neural development, and RNA interference (Corsi *et al*. 2015). Among other useful features, the availability of a large number of loss-of-function and null *C. elegans* mutants has been important for the success of *C. elegans* research. Many of these mutants have been isolated by screening worms treated with mutagens or transposons (Vallin *et al*. 2012; *C. elegans* Deletion Mutant Consortium 2012; Thompson *et al*. 2013; Kutscher and Shaham 2014). While these mutants allow us to infer the nature of a gene’s normal function and to place genes into genetic networks using epistasis analyses, many of them may also contain background mutations that can confound the interpretation of the functional importance of the gene of interest. In addition, a significant fraction of *C. elegans* genes either only have alleles that do not cause complete loss of gene function or currently do not have mutants available. As such, an efficient and convenient way to generate null mutants in the wild-type background would be of great interest. Here, we report a simple and robust method using CRISPR/Cas9 and a universal stop knock-in (STOP-IN) cassette to achieve this purpose.

The CRISPR/Cas9 system was originally identified in bacterial and archaeal immune system used to defend against viruses (Wiedenheft *et al*. 2012). Cas9 protein from the bacterium *Streptococcus pyogenes* is an RNA-guided endonuclease that creates site-specific cleavage of double stranded DNA. Cas9 protein complexes with two small RNAs to from a functional Cas9 ribonucleoprotein: a trans-activating crRNA (tracrRNA) that acts as a scaffold, and a CRISPR RNA (crRNA) that determines the target specificity of Cas9 nuclease activity (Gasiunas *et al*. 2012; Jinek *et al*. 2012). The Cas9 ribonucleoprotein recognizes the DNA target that is homologous to the first 20 nucleotides of the crRNA guide sequence and is followed by a protospacer adjacent motif (PAM), which in the case of S. *pyogenes* is 5’-NGG-3’, and generates a double-strand break three base pairs upstream of the PAM site (Figure 1A). After the break is generated, the DNA strands can be re-joined together through error-prone non-homologous end joining (NHEJ) or precise homology directed repair (HDR). The CRISPR/Cas9 system was engineered for modification of genomes in mammalian cells and has revolutionized the field of genome editing since then (Cong *et al*. 2013; Mali *et al*. 2013).

The CRISPR/Cas9 system has also been successfully adopted for genome editing in *C. elegans* by injecting plasmids, RNA and/or protein of the CRISPR/Cas9 system in the gonads of adult hermaphrodite (Chen *et al*. 2013; Cho *et al*. 2013; Katic and Großhans 2013; Lo *et al*. 2013; Tzur *et al*. 2013; Waaijers *et al*. 2013; Friedland *et al*. 2013; Dickinson *et al*. 2013; Chiu *et al*. 2013). Different strategies have been described to facilitate identification of the desired genome editing event (reviewed by Dickinson and Goldstein 2016). For example, co-conversion and co-CRISPR methods effectively enrich for animals that have the desired alternations: animals with a successful genome editing at the co-conversion locus with a dominant visible phenotype, such as *dpy-10*, are likely to have edits at another independent locus (Ward 2014; Kim *et al*. 2014; Arribere *et al*. 2014).

Several CRISPR approaches have been described to generate loss-of-function alleles or null alleles for *C. elegans* genes, in which deletions/insertions or premature nonsense mutations were introduced at specific loci. Without a repair template, error-prone NHEJ generally creates deletions or insertions with unpredicted flanking sequences in target genes after Cas9 cutting (Katic and Großhans 2013; Friedland *et al*. 2013; Chiu *et al*. 2013; Chen *et al*. 2014). In addition, some of the NHEJ-mediated insertions or deletions could be in-frame, which may only marginally reduce gene activity. For mutants that have unknown phenotypes, PCR is commonly used to detect the desired genome editing. The detection of small deletions and/or insertions via PCR is rather laborious, as the PCR products generally needs to be denatured, annealed, and treated with enzymes that recognize mismatches, such as CEL-1 nuclease and T7 Endonuclease I, to identify candidates with successful editing. When using repair templates with long homology arms, HDR following cutting by Cas9 can produce precise insertion of large exogenous sequences, such as *gfp* or an antibiotic selection cassette, to disrupt or replace the gene of interest (Chen *et al*. 2013; Tzur *et al*. 2013; Norris *et al*. 2015). Although several vectors have been developed to facilitate the preparation of such repair templates (Dickinson *et al*. 2015; Schwartz and Jorgensen 2016), laborious molecular cloning may be required to construct repair template plasmids with long homology arms. In addition, large insertions or deletions may affect the expression of the neighboring genes (Tzur *et al*. 2013). Single stranded DNA oligos with short homology arms (~30-60 nucleotides) can efficiently serve as repair templates to insert up to 130 nucleotides into the *C. elegans* genome with CRISPR (Paix *et al*. 2017). In previous attempts to making knock-outs by introducing a single stop codon, the reading frame of the target gene would need to be considered and additional mutations were also needed to prevent re-cutting by Cas9 and/or to introduce a restriction enzyme site for detection of successful genome modification (Paix *et al*. 2017).

We sought to develop an easy and efficient CRISPR/Cas9 method to generate putative null mutants with pre-defined mutations in *C. elegans*. To this end, we designed a 43-nucleotide-long universal stop knock-in (STOP-IN) cassette that will create stop codons and frameshifts in all three reading frames. We describe an efficient co-conversion CRISPR method in which Cas9 ribonucleoprotein and short single stranded DNA oligo repair templates are used to generate putative null alleles by inserting the STOP-IN cassette into the coding regions of target genes. The universal STOP-IN cassette allows the usage of a common internal primer for easy detection of putative null alleles with successful insertion via PCR. The resultant null alleles can be reverted to the original wild-type sequence via a second oligo-directed CRISPR/Cas9 editing targeting the exogenous Cas9 target site included in the STOP-IN cassette.

## Results

### Design of our universal STOP-IN cassette

We aimed to develop a cloning-free and streamlined protocol to design single stranded DNA repair oligos to generate putative mutants with well-defined mutations and to simplify the detection of knock-in alleles with PCR. To this end, we designed a 43-nucleotide-long universal stop knock-in (STOP-IN) cassette that would create putative null mutants when inserted within the exons, regardless of the reading frame. The cassette contains an exogenous Cas9 target site with a PAM site, which has been shown to function in when inserted into the *C. elegans* genome (Paix *et al*. 2017), followed by five stop codons, as well as a *Nhe*I restriction enzyme site (Figure 1B). Once inserted into the coding stand, this STOP-IN cassette will generate stop codons in three reading frames (a tandem pair of stop codons for two of the three frames and one stop codon for the third frame; Figure 1B). The universal STOP-IN cassette is 43 nucleotides and thus will also create a frameshift in the target gene even if stop codon readthrough occurs.

Insertion of similar three-frame stop cassettes has been used to knockout gene activity in zebrafish, *C. elegans* and mammalian cell lines (Gagnon *et al*. 2014; Farboud and Meyer 2015; Maruyama *et al*. 2015). The design of our STOP-IN cassette offers some unique features. The included exogenous Cas9 target site allows future re-editing of the genomic locus, and the *Nhe*I site can also be used for PCR/RFLP detection. Our cassette also allows the use of a single universal inner primer (e.g., we used the universal reverse primer P2: 5’-GCTTATCACTTAGTCACCTCTGCTC-3’) for detection of successful knock-in events, when combined with a gene-specific forward outer primer (P1 in Figure 1F). In addition, the insertion of the universal STOP-IN cassette (43 bp) is easy to identify by size shift on gel electrophoresis of a PCR product made using the outer primer set flanking the target cut site(P1 and P3 in Figure 1F), when the PCR product is less than 500 bp.

### Generation of putative *aex-2* null mutants

To test if insertion of our universal STOP-IN cassette into the coding sequence could generate putative null mutants, we chose the gene *aex-2*, which has distinctive and easily observable loss-of-function phenotypes. *aex-2* encodes a G protein-coupled receptor that is essential for rhythmic enteric muscle contraction (EMC) during the *C. elegans* defecation motor program. The strong loss-of-function *aex-2(sa3)* mutants almost completely lack EMCs, leading to a constipated phenotype with a distended intestinal lumen (Mahoney *et al*. 2008; Figures 2D-2E). We tested whether new *aex-2* alleles generated using our universal STOP-IN cassette had the same phenotype as *aex-2(sa3)* animals.

We first selected a guide sequence in the third exon of the *aex-2* gene to insert the STOP-IN cassette with CRISPR/Cas9 (Figures 2A-2B; see Supplemental File S1 for a detailed protocol). Following microinjection and identification of F_1_ animals showing the co-conversion marker (Figure 1C), 7 of the 16 F_1_ clones genotyped (44%) had successful insertion of the STOP-IN cassette (Figure S1; Table 1). Two different STOP-IN alleles of *aex-2* were retained (*sy1078* and *sy1079*), originating from different F_1_ animals, and confirmed by Sanger sequencing (data not shown). Both *sy1078* and *sy1079* animals, behaved similarly to *aex-2(sa3)* mutants and were completely defective in EMC during the defecation cycle and had distended intestine lumens (Figure 2D and 2E), consistent with the idea that both *sy1078* and *sy1079* were null alleles. Thus, our STOP-IN method can generate putative null mutants in target genes.

### Reversion of *aex-2* mutants to the wild type with a second CRISPR/Cas9

Next, we tested if we could revert the STOP-IN alleles of *aex-2* back to wild-type sequence using the exogenous Cas9 target site that was included the STOP-IN cassette. We targeted the *aex-2(sy1078)* mutants using a crRNA for the exogenous Cas9 target site, and a 71-nuceotide-long single stranded DNA repair oligo with wild-type *aex-2* sequence (Figure 2C). We obtained F_1_ animals that unlike the parental *aex-2(sy1078)* were not constipated, presumably animals with reverted alleles that showed normal EMC during the defecation cycle. We retained two revertant *aex-2* alleles (*sy1078sy1090* and *sy1078sy1091*) and confirmed the wild-type sequence of the *aex-2* locus around the Cas9 target site with Sanger sequencing.

Both *aex-2* revertants showed wild-type level of EMC and had normal intestinal lumens (Figures 2E-2F). Thus, the null alleles with the STOP-IN cassette can be reverted back to the wild type. The reversion of a null mutant to the wild type offers a convenient and convincing way to prove that the phenotype associated with a null mutation is caused by loss of the gene’s function.

### Calponin-like protein knockouts have pharyngeal pumping defects

We used our STOP-IN method to generate null alleles for the poorly understood and uncharacterized *clik* genes. These genes have been shown to be highly expressed and almost completely specific for pharyngeal muscle: *clik-1, clik-2*, and *clik-3* (Yuet *et al*. 2015). These genes encode proteins that share some homology with human Calponin-1 (CNN-1), a protein found in smooth muscle cells. Two unique types of protein domains are found in CNN-1: a single calponin homology (CH) domain, and multiple calponin family repeats (CFRs). The three *C. elegans* genes we chose all contain varying numbers of CFRs (Figure 3A). CFRs are thought to be actin-interacting domains, negatively affecting actin-cofilin interactions (Yamashiro *et al*. 2007). Given the highly specific expression of the *clik* genes in the pharynx, we generated putative null mutants of these genes and assayed for pumping defects.

**Figure 3.**
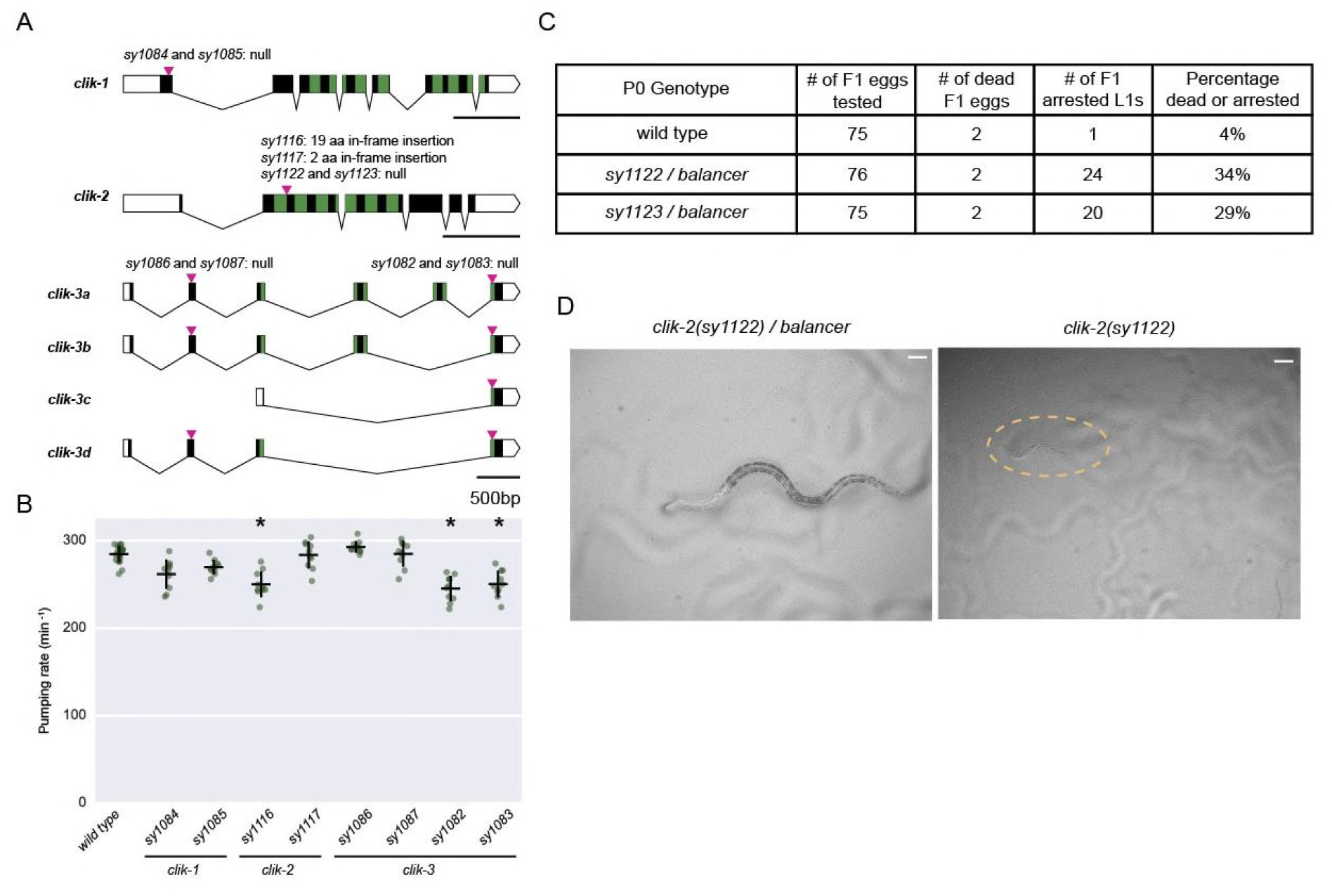
Mutants of *clik* genes and pharyngeal pumping. (**A**) Gene structures of three *C. elegans* genes encoding Calponin-like proteins: *clik-1, clik-2*, and *clik-3*. Green blocks represent the calponin family repeats (CFRs). Magenta arrowheads indicate Cas9 cut sites where the STOP-IN cassette inserted in indicated alleles, except for *sy1116* and *sy1117*, which are small insertions likely resulted from NHEJ during repair. aa, amino-acid. Scale bars, 500 bp. (**B**) Quantification of pumping rate of *clik-1, clik-2, clik-3* mutants. Each green dot represents the mean pumping rate of one adult animals. The *clik-2(sy1116)* partial loss-of-function mutants and *clik-3(sy1082)* and *clik-3(sy1083)* null mutants have reduced pumping rates, compare to the wild type. Horizontal bars are the mean, vertical bars are SEM. n = 10 for all genotypes. * P < 0.05, One-way ANOVA with Tukey’s post-test. (**C**) Table showing that *clik-2(sy1122)* and *clik-2(sy1123)* null mutants are recessive lethal. L1, the first larval stage of *C. elegans*. The *balancer* allele used in (C and D) is *tmC30[tmIs1243]*, an aneuploidy-free and structurally defined balancer for the part of chromosome X where *clik-2* is located. (**D**) Bright field images showing *clik-2(sy1122)/balancer* heterozygote (left) and *clik-2(sy1122)* homozygote (right), roughly 16-19 hours after hatching. Scale bars are 20 μm.

We inserted the STOP-IN cassette in early exons for *clik-1* and *clik-2*, both of which have single isoforms (Figure 3A). *clik-3* has four isoforms, of which the c isoform is the shortest and consists of the last exon. Thus, we created two different alleles of *clik-3* in which the STOP-IN cassette was inserted at different locations: one in the second exon (*sy1086* and *sy1087*, affecting isoforms a, b, and d) and the other in the last exon (*sy1082* and *sy1083*, affecting all isoforms, Figure 3A).

We used these alleles to determine if any of these three genes are involved in the regulation of pharyngeal pumping. *clik-1* null mutants (*sy1084* and *sy1085*) were similar to the wild type, suggesting that *clik-1* is not essential for pharyngeal pumping (Figure 3B). For *clik*-3, we found that only the *sy1082* and *sy1083* alleles affecting all isoforms displayed a mild (14% decrease) pumping defect. The *sy1086* and *sy1087* alleles affecting isoforms a, b, and d, did not exhibit pumping defects (Figure 3B).

We were unable to homozygose the *clik-2* null alleles with the STOP-IN cassette, *sy1122* and *sy1123*, leading us to believe *clik*-2 null mutants were lethal. We thus created balanced *sy1122/tmC30[tmIs1243]* and *sy1123/tmC30[tmIs1243]* strains using a multiple-inversion balancer (Dejima et al., 2018). These animals appeared wild-type for growth and pumping. However, their progeny included tiny, starved L1-arrested animals which did not appear to pump, in roughly Mendelian ratios (Figures 3C-3D). Close observation of these arrested animals showed barely perceptible fasciculation of pharyngeal muscles and the grinder, but no full contractions. When constructing our *clik-2* null mutants, we also obtained two inframe insertion alleles for *clik-2, sy1116* (19-amino-acid insertion) and *sy1117* (2-amino-acid insertion), which were likely to be resulted from error-prone NHEJ. Pharyngeal pumping rate was decreased by about 15% in *sy1116* animals; *sy1117* animals appeared normal (Figure 3B).

### Robustness of the knock-in method

To test the robustness of our STOP-IN method to generate putative null mutants, we used it to generate putative null mutants for additional 17 genes, most of which did not have null mutants available. We found that, on average, 46% of the F_1_ animals with co-conversion marker (Rol for *dpy-10* or Unc for *unc-58*) we screened (n = 694) had the successful insertion of the STOP-IN cassette (ranging from 11% to 100%, Table 1). We noticed that what we could not homozygose some of the alleles we obtained, and we speculated that these mutants were likely to be lethal (Table 1) Note that the knock-in efficiency was markedly lower in those putative lethal genes, than for non-lethal genes (27 % vs 59%; n = 267 and 427, respectively, P < 0.0001, Chi-square test). However, the knock-in efficiency for the lethal genes may be underestimated, as we could not recover the F_1_ animals for these genes that contained two loss-of-function alleles induced by CRISPR/Cas9.

While most of the candidate F_1_ animals with desired knock-in events were heterozygotes, we noticed that for some genes, a small fraction of F_1_ animals appeared to be homozygous for the knock-in null alleles, as they had a correct PCR product with the outer and the internal primer pair, but only had the bigger PCR band for the knock-in allele with the outer primer pair (Examples shown in Figures S1 and S2). However, it was also possible that these putative F_1_ homozygotes could be heterozygotes. e.g., one successful knock-in allele and the other allele with a deletion that removed a region where one of both external primers bind. Indeed, CRISPR has been shown to create two independent heritable alleles in a single F_1_ animal (Cho *et al*. 2013). As we generally had more than two F_1_ animals that are heterozygotes and to make sure the homozygotes were from single alleles, we always singled out several F_2_ progeny from each of independent heterozygote F_1_ animals and identified the homozygotes for null knock-in alleles by genotyping the F_3_ progeny.

We found that partial repair occurred in a small fraction of animals that we screened, in which the knock-in PCR band with the outer primer pair appeared to be slightly smaller than the predicted size (Figures S1 and S2). Such partial repair has been reported before (Arribere *et al*. 2014). Thus, we recommended validating homozygote knock-ins by sequencing the PCR products with the outer primer pairs in F_3_ homozygotes.

## Discussion

We report a simple and robust CRISPR/Cas9 method to generate putative *C. elegans* null mutants by inserting a 43-neucleotide-long stop knock-in (STOP-IN) cassette in exons of target genes. We report high knock-in efficiency (46%, n = 694) in F_1_ co-converted animals that we screened. We applied our STOP-IN method to generate null mutants for 21 genes, including three *clik* genes which are specifically expressed in the pharyngeal muscles (Yuet *et al*. 2015).

We showed that *clik-2* was essential for pumping in *C. elegans*. As a proof of concept, we also showed that null alleles generated in our STOP-IN method can be reverted to wild type by a second genome editing targeting the exogenous target site included in the STOP-IN cassette.

### Useful features of our STOP-IN method to generate null mutants

The main features of STOP-IN method are as follows. First, it is straightforward to design the guide sequence and the repair template, without the need to consider the reading frame of the target gene at the engineered locus and to introduce other mutations to create a restriction enzyme or primer binding site for detection of a successful insertion. Second, similar to previously described CRISPR/Cas9 protocols that used preassembled Cas9 ribonucleoprotein (Cho *et al*. 2013; Paix *et al*. 2015), our method is cloning-free, as all the reagents for the CRISPR experiments, including Cas9 protein, tracrRNA, crRNAs, and single stranded DNA repair oligos, are commercially available. Third, our method is highly efficient. 46% of F_1_ animals with a co-conversion phenotype that we genotyped had the desired knock-in events. Fourth, combined with an outer primer specific to the gene of interest, the common inner primer designed in the universal STOP-IN cassette makes it easy to detect successful knock-in events by PCR. Fifth, the null alleles generated with our method can be reverted to the wild type allele with a second genome editing on the exogenous Cas9 target sequence that is included in the STOP-IN cassette and supplemented with the oligo containing the original *C. elegans* genome sequence. This can be used to prove that the null mutation in the gene of interest is causative for the phenotype observed. In principle, the exogenous Cas9 target site can be also used to other types of genome editing. For example, inserting an in-frame *gfp* reporter fused to the gene of interest.

### Potential modifications of our CRISPR method

The modular design of the STOP-IN cassette offers opportunities for future modification. While we selected the exogenous Cas9 target site with a PAM site (5’-GGGAAGTTTGTCCAGAGCAG<u>AGG</u>-3’) and the restriction enzyme NheI (5’-gctagc-3’) in our current design, it can be modified to generate null alleles, as long as the core sequence with the 3-frame stop codons and frameshifts is preserved. For example, the exogenous Cas9 target site can be replaced if an improved sequence is reported for *C. elegans*. Similarly, the restriction enzyme site can be changed to others, as desired.

In this study, we successfully used *dpy-10* or *unc-58* as co-conversion markers to enhance detection of CRISPR knock-in events for our STOP-IN method. Other reported co-conversion markers, such as *pha-1, unc-22, sqt-1* and *rol-6* (Ward 2014; Kim *et al*. 2014; Arribere *et al*. 2014), should work as well. In principle, our STOP-IN method should also work with other CRISPR/Cas9 delivery strategies, including injecting mRNA or plasmids that express Cas9 and guide RNAs (reviewed by Dickinson and Goldstein 2016), instead of pre-assembled Cas9 ribonucleoprotein complex used in this study. While these alternative strategies may make our STOP-IN method more useful in other experimental systems, we expect that the knock-in efficiency with injection of mRNA or plasmids may be lower than delivery of pre-assembled Cas9 ribonucleoprotein (Farboud *et al*. 2018).

Unlike deletions obtained from CRISPR/Cas9 protocols not involving HDR, which generally have unpredictable endpoints and may affect gene activity in unknown ways, null alleles generated using our STOP-IN method are well defined and reproducible: the alleles of the target gene generated using the same guide RNA and the repair template will have the exactly same genetic mutation. This consistency will increase robustness in comparing interactions between genes. With its high efficiency, our STOP-IN method can be used to generate molecularly indistinguishable null alleles of the target gene in multiple genetic backgrounds: wild-type background, existing mutants, or in other isolates of the same species.

In this way, one can compare single, double, and even multiple mutants to infer genetic interactions and examine complex effects of the target gene in different genetic backgrounds. Our method is particularly useful when close linkage impedes creation of double-mutant animals by traditional recombination. In addition, as the relative small universal STOP-IN cassette is likely to have minimal effects on gene structure, guide sequences in isoform-specific exons may be used to generate isoform-specific null mutations, as we did for *clik-3* (Figure 3A). With its easy design and high knock-in efficiency, we envision that our universal STOP-IN cassette will facilitate the generation of new null mutants in *C. elegans* and potentially other genetic model organisms, which can be used to understand gene function.

## Acknowledgments

We thank Tsui-Fen Chou for providing Cas9 protein and Shohei Mitani for the balancer strain FX30236. We thank Barbara Meyer for sharing unpublished results and Hillel Schwartz for discussion. We also thank the members from the Sternberg lab for comments on the manuscript. Some strains were provided by the CGC, which is funded by NIH Office of Research Infrastructure Programs (P40 OD010440). WormBase annotations were crucial for choosing target genes and location of guide sequences. This work was supported NIH (K99GM126137 to H.W., R24OD023041 to P.W.S., and T32GM007616, which supported J.L.).

---

**Figure S1.**
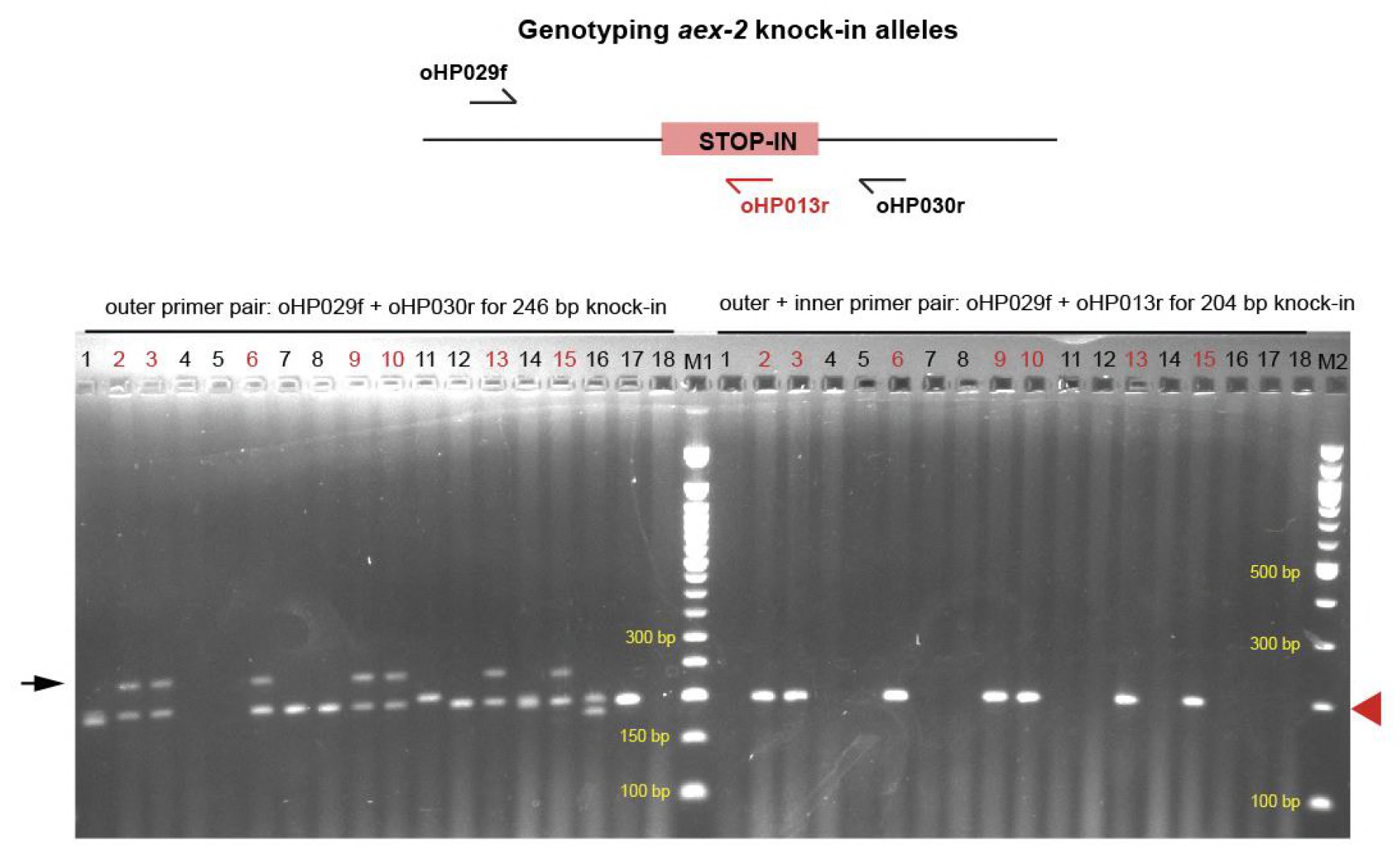
Genotyping result to identify null *aex-2* alleles with the universal STOP-IN cassette. (**A**) Diagram of the primers used for genotyping *aex-2*. The outer primer set (oHP029f and oHP030r) will yield a 246 bp PCR product from the knock-in allele and a 203 bp PCR product from wild type allele; the outer and inner primer set (oHP029f and oHP013r) will produce a 204 bp PCR product from the knock-in allele and no PCR product from wild type allele. The red rectangle represents the universal STOP-IN cassette described in the manuscript. (**B**) Electrophoresis of PCR products from the genotyping experiments with indicated primers of F_2_ progeny from 16 (from 1 to 16) different F_1_ animals that had the co-conversion phenotype, Rol. 17 and 18 were controls: the wild-type strain used for injection and the bacterial food for *C. elegans*, respectively. The black arrow on the left indicates the correct PCR band (246 bp) from the desired knock-in alleles from the PCR reaction using oHP029f and oHP030r. The red arrowhead on the right indicates the correct band (204 bp) for the desired knock-in alleles from the PCR reaction using oHP029f and oHP013r. The samples with numbers in red (2, 3, 6, 9, 10, 13, and 15) were heterozygous for the knock-in alleles. Samples 1 and 16 were heterozygous a small deletion. Sample 11 appeared to be homozygous for a small insertion. The PCR reactions appeared to fail in samples 4 and 5. M1 and M2 represent 50 bp and 100 bp DNA ladders, respectively (NEB, Ipswich, MA).

**Figure S2.**
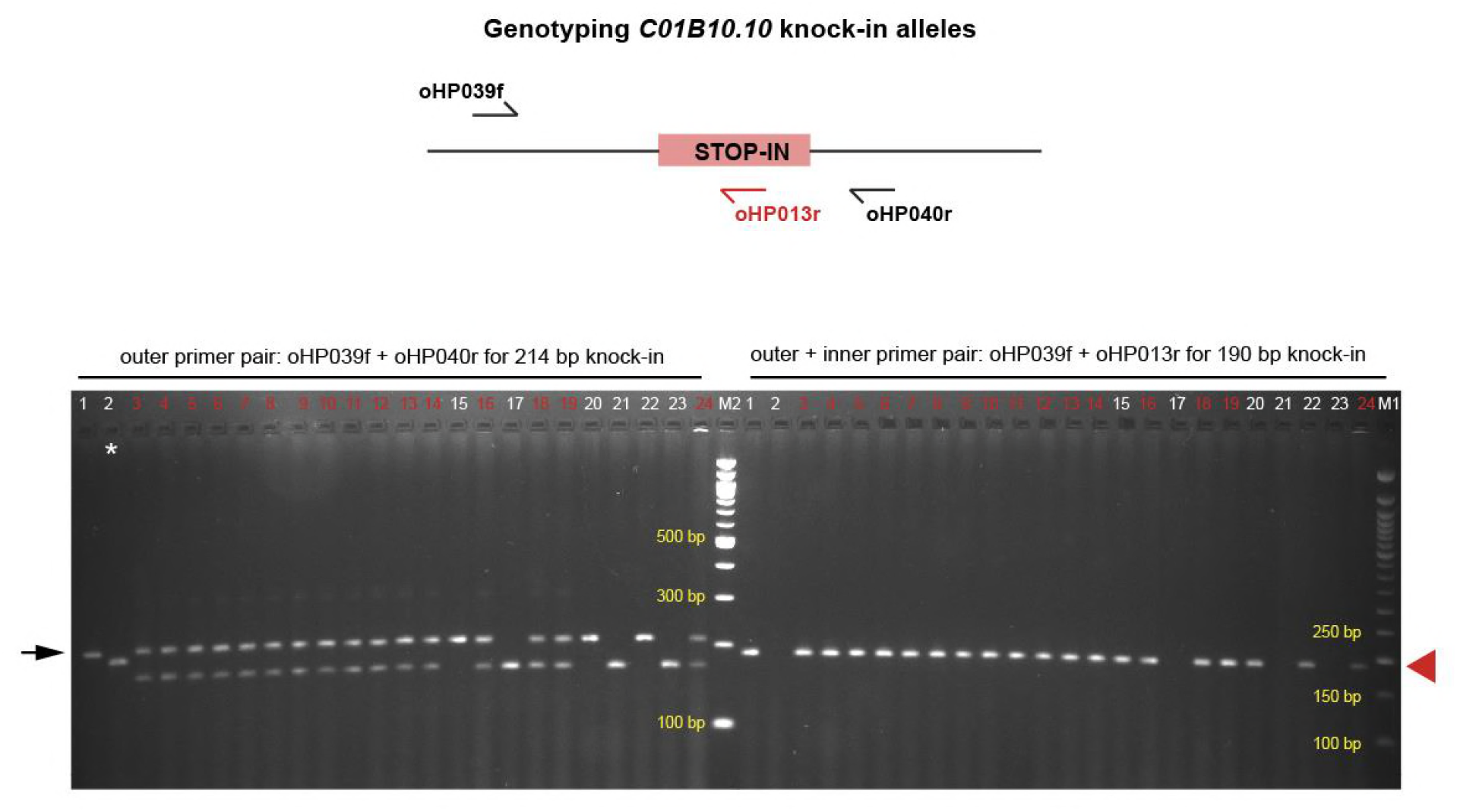
Genotyping result to identify null *C01B10.10* alleles with the universal STOP-IN cassette. (**A**) Diagram of the primers used for genotyping *C01B10.10*. The outer primer set (oHP039f and oHP040r) would yield a 214 bp PCR product from the knock-in allele and a PCR 171bp product from the wild type allele; the outer and inner primer set (oHP039f and oHP013r) would only produce a PCR product of 190 bp band for the knock-in allele and no product from the wild type allele. The red rectangle represents the STOP-IN cassette described in the manuscript. (**B**) Electrophoresis of PCR products with indicated primers from genotyping F_2_ progeny of 24 different F_1_ animals that had the co-conversion phenotype, Rol. The black arrow on the left indicates the correct band (214 bp) for successful knock-in alleles from the PCR reaction using oHP039f and oHP040r. The red arrowhead on the right indicates the correct band (190 bp) for successful knock-in allele from the PCR reaction with oHP39f and oHP13r. The 16 samples in red (3, 4, 5, 6, 7, 8, 9, 10, 11, 12, 13, 14, 16, 18, 19 and 24) were heterozygotes for the desired knock-in alleles. Four samples (1, 15, 20, and 22) were putative homozygotes for the desired knock-in allele. Sample 2 was likely to homozygous for a small insertion. The three samples left (17, 21, and 23) were likely to be wild-type alleles. M1 and M2 represent 50 bp and 100 bp DNA ladders, respectively (NEB, Ipswich, MA).

